# Admixture mapping in interspecific *Populus* hybrids identifies classes of genomic architectures for phytochemical, morphological and growth traits

**DOI:** 10.1101/497685

**Authors:** Luisa Bresadola, Céline Caseys, Stefano Castiglione, C. Alex Buerkle, Daniel Wegmann, Christian Lexer

## Abstract

- The genomic architecture of functionally important traits is key to understanding the maintenance of reproductive barriers and trait differences when divergent populations or species hybridize. We conducted a Genome-Wide Association Study (GWAS) to study trait architecture in natural hybrids of two ecologically divergent *Populus* species.
- We genotyped 472 seedlings from a natural hybrid zone of *Populus alba* and *P. tremula* for genome-wide markers from reduced representation sequencing, phenotyped the plants in common gardens for 46 phytochemical (phenylpropanoid), morphological, and growth traits, and used a Bayesian polygenic model for mapping.
- We detected three classes of genomic architectures: (1) traits with finite, detectable associations of genetic loci with phenotypic variation in addition to highly polygenic heritability, (2) traits with indications for polygenic heritability only, (3) traits with no detectable heritability. For class (1), we identified genome regions with plausible candidate genes for phenylpropanoid biosynthesis or its regulation, including MYB transcription factors and glycosyl transferases.
- GWAS in natural, recombinant hybrids represents a promising step towards resolving the genomic architecture of phenotypic traits in long-lived species. This facilitates the fine-mapping and subsequent functional characterization of genes and networks causing differences in hybrid performance and fitness.

## Introduction

Understanding the genetic architecture of phenotypic trait differences between divergent populations and species has long been a fundamental goal in evolutionary genetics. At the within-species level, interest has primarily been on understanding local adaptation in wild species and the selective forces operating during domestication of agriculturally important species (Atwell *et al*., 2010; Huang *et al*., 2011; Jones *et al*., 2012; Li *et al*., 2012; Evans *et al*., 2014). At the between-species level, evolutionary geneticists have sought to understand the origin and maintenance of adaptive trait differences and reproductive barriers between species, and thus the mechanisms maintaining species integrity (Coyne & Orr, 2004; Feder *et al*., 2012; Lindtke *et al*., 2013; Turner & Harr, 2014).

The genetic architecture of traits is frequently inferred from family-based association studies or experimental crosses (e.g. Tanksley, 1993; Kong *et al*., 2013; Liller *et al*., 2017), but both methods are limited to organisms that exhibit short generation times, are readily crossed in the greenhouse or laboratory, and produce abundant offspring to yield sufficient power for mapping (e.g. Bradshaw *et al*., 1998; Rieseberg *et al*., 2003; Zhu *et al*., 2003). One way to extend genetic mapping to longer-lived organisms is to use recombinants from natural hybrid zones (Barton & Hewitt, 1985; Rieseberg & Buerkle, 2002; Buerkle & Lexer, 2008). However, admixture mapping, which was originally introduced by human geneticists (Chakraborty & Weiss, 1988), has only rarely been applied to plant and animal species (sunflowers, Rieseberg *et al*., 1999; sticklebacks, Malek *et al*., 2012; poplars, Lindtke *et al*., 2013 and Suarez-Gonzalez *et al*., 2018; canids, vonHoldt *et al*., 2016; warblers, Brelsford *et al*., 2017).

What makes admixture mapping attractive is the opportunity to analyze trait differences segregating in natural populations. Compared to mapping in controlled crosses, much more of the phenotypic and genetic variation of wild species may be captured (Lexer *et al*., 2004; Buerkle & Lexer, 2008). A challenging aspect, however, is the complexity of Linkage Disequilibrium (LD) along the genome, which may be affected by genomic incompatibilities and coupling effects expected in hybrid zones of highly divergent populations (Barton & Hewitt, 1985; Bierne *et al*., 2011; Lindtke *et al*., 2013; Gompert *et al*., 2017).

Hybrid zones formed by *Populus* species represent textbook examples of natural interspecific crosses (Stettler *et al*., 1996; Arnold & Kunte, 2017). *Populus* is a model genus for studies of tree form, function, and evolution of forest foundation species, including their involvement in eco-evolutionary dynamics (Tuskan *et al*., 2006; Whitham *et al*., 2006). This study is focused on the ecologically divergent *Populus alba* (White poplar), widespread in Southern Eurasia and Northern Africa, and *P. tremula* (European aspen), found mainly in Northern Eurasia. Even after >2.8 million years of divergence (Christe *et al*., 2017), the reproductive barriers between these species are incomplete, thus they hybridize in regions where their ranges overlap (Christe *et al*., 2016; Macaya-Sanz *et al*., 2016; Zeng *et al*., 2016). Despite strong postzygotic barriers, a broad range of recombinant hybrid seeds are formed in these hybrid zones (Lindtke *et al*., 2014; Christe *et al*., 2016). Here, we use a recombinant mapping population composed of plants grown from seeds collected from open pollinated trees in a natural hybrid zone and cultivated in two common gardens. Using these, we study a range of functionally and ecologically relevant traits exhibiting phenotypic differentiation among the parental species and their hybrids, including phytochemical traits (the abundances of phenylpropanoid secondary metabolites in leaves), leaf morphology, and growth-related characters.

Recent results from Genome-Wide Association Studies (GWASs) in trees, humans, and other species suggest that the architecture of adaptive traits is often polygenic, that is, they are not determined by a few genes of large effect, but rather by many loci with small effect (Pritchard *et al*., 2010; Rockman, 2012; Evans *et al*., 2014; Pasaniuc & Price, 2017). Our results highlight the importance of both finite, sparse genetic loci and highly polygenic heritability of quantitative traits. Knowing these contributions is important for any in-depth molecular genetic or genomic study aimed at dissecting complex, functionally important traits. For those traits for which “finite” architectures were detected, we identified candidate genes located within associated genomic regions and discuss potential molecular mechanisms underlying trait variation.

## Materials and Methods

### Plant materials

We analyzed seedlings of *P. alba* L., *P. tremula* L., and their hybrids, a.k.a. *P. x canescens (Aiton) Sm.* All seeds were collected from open-pollinated mother trees in the Parco Lombardo della Valle del Ticino in the North of Italy (Lexer *et al*., 2010; Lindtke *et al*., 2014). Seeds were sampled in 2010, 2011 and 2014 and germinated broadly following Lindtke *et al.* (2014). Approximately two months after germination, we moved seedlings to larger pots and arranged them in a common garden using a block design with randomized positions within blocks. We grew ca. 500 seedlings from 39 families in two locations: at the Botanical Garden of the University of Fribourg, Switzerland and at the University of Salerno, Italy.

### Genetic data

We conducted a Restriction site Associated DNA sequencing (RAD-seq) experiment as follows: First, we extracted DNA from silica-dried leaf material of 472 individuals using the Qiagen DNeasy Plant Mini Kit (Valencia, CA) and standardized concentrations to 20 ng/μl. Second, we submitted all samples to Floragenex (Eugene, OR), where five libraries with 95 individuals each were prepared according to their standard commercial procedure, very similar to the original RAD-seq protocol (Baird *et al*., 2008). Specifically, genomic DNA was digested with the restriction enzyme *PstI*, chosen according to previous RAD-seq studies of these species and their hybrids (Stölting *et al*., 2013; Christe *et al*., 2016), and the libraries were sequenced single-ended on one lane of an Illumina HiSeq2500 instrument each (SRA accession number XXXX). Third, we processed RAD-seq data using state-of-the-art tools, including mapping to the *P. trichocarpa* reference genome with Bowtie2 2.2.4 (Langmead & Salzberg, 2012) and variant calling with GATK 3.4.46 (DePristo *et al*., 2011) following best practice (Supporting Information Methods S1). Fourth, we applied strict filters to only retain reliable sites: we removed all sites with more than two segregating alleles or with an average depth above the 95% quantile to exclude potentially paralogous loci. To avoid Single Nucleotide Variants (SNVs) originating from misalignments, we further removed indels and variant sites within 5 bp of all indels that we could identify confidently using GATK on the full data.

### Inference of genome-wide ancestry

We estimated genome-wide ancestry *(q)* using entropy (Gompert *et al*., 2014) directly from genotype likelihoods after removing SNVs with minor allele frequency <0.05 or >50 % missing data and after correcting genotype likelihoods for biases associated with RAD-seq (Methods S2). We further calculated F_ST_ between parental species using Hudson’s estimator (Hudson *et al*., 1992) on the parental allele frequencies inferred with entropy.

### Inference of local ancestry

Despite substantial genetic differentiation between the parental species, they share alleles and genotypes at many loci, so that LD in our study population decays rapidly with physical distance. Thus, we used local ancestry for trait mapping, which provided greater LD along chromosomes (see below). LD is required for mapping because causal allelic variants are unlikely directly observed in reduced representation studies. We estimated local ancestry using RASPberry (Wegmann *et al*., 2011), which implements a Hidden Markov Model (HMM) to explain haplotypes of hybrid individuals as a mosaic of reference haplotypes provided for each species.

We obtained reference haplotypes by phasing previously characterized pure *P. alba* and *P. tremula* individuals (51 each) from the Italian, Austrian and Hungarian hybrid zones (Christe *et al*., 2016) using FastPhase (Scheet & Stephens, 2006), building input files with fcGENE (Roshyara & Scholz, 2014). For use in RASPberry, individuals in the reference panels were not allowed to have missing data. The genotype calling step in our common garden individuals was therefore restricted to the 45,193 SNVs covered in all parental individuals (Christe *et al*., 2016). We further masked all genotype calls based on less than five reads.

To infer local ancestries with RASPberry we used the mutation rates previously estimated for *P. alba* and *P. tremula* (Christe *et al*., 2016). As a prior on the switching probabilities we further used 5 cM/Mb as the default recombination rate, as estimated for *P. trichocarpa* (Tuskan *et al*., 2006), and sample-specific genome-wide ancestry *q* estimated using ADMIXTURE (Alexander *et al*., 2009) on all 472 individuals jointly. The remaining parameter settings, initial optimization runs, and incorporation of RAD-seq genotyping error rates are described in detail in Methods S2.

For mapping, we then used the expected ancestry genotype calculated from the posterior probabilities obtained with RASPberry. We verified the presence of LD by calculating the pairwise squared correlation between point estimates of local ancestries and visualized the results using the package LDheatmap (Shin *et al*., 2006) in R (R Core Team, 2016).

### Phenotypic data

We used 46 phenotypes, classified into phytochemical, morphological and growth traits (Table 1). For phytochemical traits, we focused on secondary metabolites in leaves from three different branches of the phenylpropanoid pathway: chlorogenic acids, salicinoids and flavonoids (see Methods S3 for more rationale on trait choice). These secondary metabolites were previously quantified for a subset of 133 samples using Ultra-High-Pressure-Liquid-Chromatography Quadrupole-Time-Of-Flight Mass-Spectrometry (Caseys *et al*., 2012, 2015) and we completed this measurement here for all 266 samples germinated in 2011.

**Table 1.** List of phenotypic traits analyzed in this study.

| Category | Trait | Abbr. | n<sup>a</sup> | Cov.<sup>b</sup> | Binary<sup>c</sup> |
| --- | --- | --- | --- | --- | --- |
| Phytochemical,<br>chlorogenic acid | 3-Caffeoyl quinic acid | C1 | 266 | q, cg | no |
|  | 3-Coumaroyl quinic acid | C2 | 266 | q, cg | no |
|  | 5-Caffeoyl quinic acid | C3 | 266 | q, cg | no |
|  | 3-Feruloyl quinic acid | C4 | 266 | q, cg | no |
|  | 1-Caffeoyl quinic acid | C5 | 266 | q, cg | no |
|  | 5-Coumaroyl quinic acid | C6 | 266 | q, cg | no |
|  | Coumaroyl quinic acid isomer | C6b | 133 | q | no |
|  | (1,5) Dicafeoyl quinic acid | C7 | 266 | q, cg | no |
| Phytochemical,<br>salicinoid | Salicin | C8 | 266 | q, cg | no |
|  | Salicortin | C9 | 266 | q, cg | no |
|  | Salicortin isomer 1 | C9i | 266 | q, cg | no |
|  | Salicortin isomer 2 | C9ii | 133 | q | no |
|  | Salicortin isomer 3 | C9iii | 266 | q, cg | no |
|  | Acetyl-salicortin | C10 | 266 | q, cg | no |
|  | Acetyl-salicortin isomer 1 | C10i | 266 | q, cg | no |
|  | Acetyl-salicortin isomer 2 | C10ii | 266 | q, cg | no |
|  | HCH-salicortin | C12 | 266 | q, cg | no |
|  | Tremuloidin | C13 | 266 | q, cg | no |
|  | Tremulacin | C14 | 266 | q, cg | no |
|  | Tremulacin isomer | C14i | 266 | q, cg | no |
|  | HCH-tremulacin | C15 | 266 | q, cg | no |
|  | Acetyl-tremulacin | C16 | 266 | q, cg | yes |
| Phytochemical,<br>flavonoid | Catechin | C17 | 266 | q, cg | no |
|  | Quercetin-rutinoside-pentose | C18 | 266 | q, cg | no |
|  | Quercetin-glucuronide-pentose | C19 | 266 | q, cg | yes |
|  | Quercetin-hexose-pentose | C20 | 266 | q, cg | no |
|  | Kaempferol-rutinoside-pentose | C21 | 266 | q, cg | yes |
|  | Isorhamnetin-rutinoside-pentose | C22 | 266 | q, cg | yes |
|  | Quercetin-3-O-rutinoside | C23 | 266 | q, cg | yes |
|  | Quercetin-3-O-glucuronide | C24 | 266 | q, cg | no |
|  | Quercetin-3-O-glucoside | C25 | 266 | q, cg | no |
|  | Kaempferol-3-O-rutinoside | C26 | 266 | q, cg | yes |
|  | Isorhamnetin-3-O-rutinoside | C27 | 266 | q, cg | yes |
|  | Quercetin-3-O-arabinopyranoside | C28 | 266 | q, cg | yes |
|  | Kaempferol-glycuronide | C29 | 266 | q, cg | yes |
|  | Quercetin-rhamnoside | C30 | 266 | q, cg | no |
|  | Isorhamnetin-glycoside | C31 | 266 | q, cg | no |
|  | Isorhamnetin-glycuronide | C32 | 266 | q, cg | yes |
|  | Isorhamnetin-acetyl-hexose | C33 | 266 | q, cg | yes |
|  | Isorhamnetin-rhamnoside | C34 | 266 | q, cg | yes |
| Morphological | Leaf area | LFAREA | 445 | q, cg, y | - |
|  | Leaf shape | LFSHAP | 445 | q, cg, y | - |
| Growth | Height first year | HEIGHT1 | 321 | q, y | - |
|  | Height second year | HEIGHT2 | 258 | q, cg | - |
|  | Diameter first year | DIAM1 | 323 | q, y | - |
|  | Diameter second year | DIAM2 | 258 | q, cg | - |

<sup>a</sup>Number of individuals with trait data
<sup>b</sup>Covariates included: *q* = genome-wide ancestry, *cg* = common garden location, *y* = planting year
<sup>c</sup>Whether the presence or absence of the chemical compound was also mapped as a binary trait in GEMMA.

For morphological traits, we measured the leaf area (LFAREA) and leaf shape (LFSHAP), known to be strongly divergent between *P. alba* and *P. tremula* (Lexer *et al*., 2009). To account for within-individual variability, we followed Lindtke *et al.* (2013) and (1) measured four leaves per plant using a ruler with a precision of 1 mm, (2) averaged the lengths and widths for each seedling, and (3) calculated LFAREA from these. LFSHAP was calculated dividing the average leaf length by the average leaf width (Lindtke *et al*., 2013).

For growth traits we included measures of height and diameter of the seedlings at one and two years after planting (HEIGHT1, HEIGHT2, DIAM1 and DIAM2). Height was quantified with a tape measure from the soil to the top of the main stem with a precision of 1 cm, while diameter was assessed with calipers with a precision of 1 mm at 10 cm above the soil. HEIGHT1 and DIAM1 were available for seedlings planted in 2010, seedlings planted in 2011 in Fribourg (not in Salerno) and seedlings planted in 2014. HEIGHT2 and DIAM2 were available only for seedlings planted in 2011, both in Fribourg and Salerno. To examine how phenotypic variation relates to *q*, we quantified the proportion of phenotypic variance explained by this variable using linear regressions. We then bootstrapped the data 1,000 times to obtain confidence intervals.

### Admixture mapping

To carry out GWAS by admixture mapping, we used the Bayesian Sparse Linear Mixed Model (BSLMM; Zhou & Stephens, 2012) in GEMMA 0.94.1 (Zhou *et al*., 2013) because it implements a polygenic approach, in which the effects of multiple loci on the phenotype are evaluated simultaneously. This provides a more complete view of genomic architecture than simpler linear models and avoids large numbers of significance tests. The set of parameters estimated by GEMMA includes *PVE* (the proportion of phenotypic variance explained by the sparse effects and random effects estimated from the kinship matrix), *PGE* (the proportion of *PVE* explained by the sparse effects only) and *n_gamma* (the putative number of sparse effect loci involved in determining the phenotype). The product of *PVE* and *PGE* gives the proportion of total phenotypic variance explained by sparse effects, which is commonly referred to as narrow-sense heritability *h^2^*.

GEMMA also estimates the probability of each locus to have a detectable sparse effect on the phenotype (the Posterior Inclusion Probability, PIP). To aggregate information from neighboring SNVs along the genome, we summed PIPs over windows of 0.5 Mb as we found large windows to result in similar patterns but making the identification of candidate genes harder (Fig. **S1**). We selected windows with a PIP ≥0.4 for further analysis of candidate genes, which is a higher threshold compared to other studies (Gompert *et al*., 2013; Comeault *et al*., 2014; Chaves *et al*., 2016).

To account for non-independence among samples, and to attribute phenotypic variation to overall genetic composition of individuals (the highly polygenic component of heritability), GEMMA estimates a kinship matrix from the genetic data and includes it as covariate in the mixed model. Since we used local ancestries as genetic input, this kinship matrix is effectively a genomic similarity matrix and captures differences in ancestry across individuals and families (Fig. **S2**). Prior to running GEMMA, we further regressed out q, the planting year and the common garden location from the phenotypes using a linear model to account for their potentially confounding effects.

For each trait, we ran 10 independent Markov chains of 12 million iterations and discarded the first 2 million as burn-in. To evaluate the robustness of our conclusions, we also ran GEMMA including the covariates in the input file (-*notsnp* option), rather than regressing them out. For the 12 phytochemical compounds with zero abundance in >10% of individuals, we further coded trait values as presence (1) and absence (0) and used binomial logistic regression to obtain residuals used as phenotypic information. Additional information on GEMMA models and parameter settings can be found in Methods S4.

#### Analysis of traits with accessible, sparse genomic architecture

We selected a core set of traits with evidence for sparse, finite architectures for further analysis. These traits had an estimated *h^2^* ≥0.01 and *n_gamma* >0 with at least 95% posterior probability. For these traits we then selected windows with PIP ≥0.4 (also for models of binary traits) and retrieved genes annotated in them in the *P. trichocarpa* reference genome (Ptrichocarpa_210_v3.0; Tuskan *et al*., 2006) and in *Arabidopsis thaliana* (The Arabidopsis Information Resource (TAIR); Berardini *et al*., 2015) to identify orthologous genes. We then examined the list of genes for candidates putatively involved in the control and modulation of the phenotypes analyzed in this study.

## Results

### Genome-wide ancestry

After filtering, we kept 127,322 SNVs to infer genome-wide ancestry (*q*) across individuals. We found considerable variation in the genomic composition of the seedlings, spanning the full range between the parental species (0 and 1; Fig. **1a**). The average F_ST_ between the species was 0.3922, which is very similar to previous estimates based on a range of different molecular data (Lexer *et al*., 2007; Stölting *et al*., 2013; Christe *et al*., 2016). Despite this elevated differentiation, most alleles were shared between species, with only 11.6% showing allele frequency difference >0.95.

**Fig. 1.**
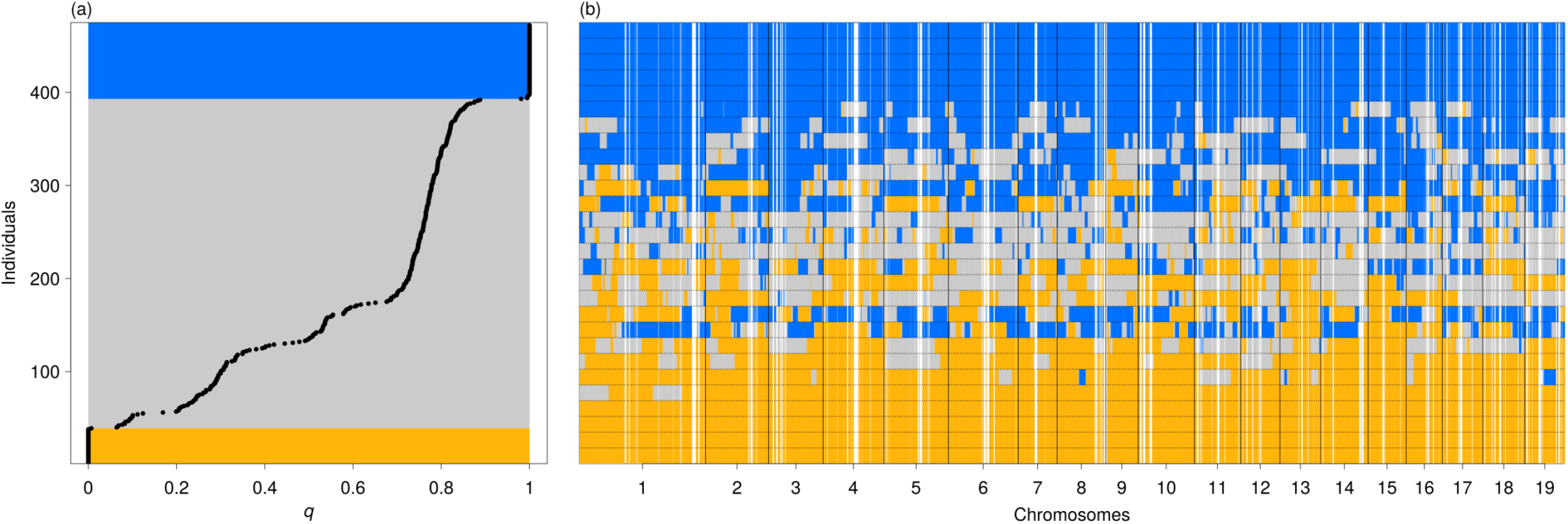
Genomic composition of Genome-Wide Association Study (GWAS) panel. (**a**) Genome-wide ancestry *q* for each common garden seedling, as estimated by entropy. Orange and blue rectangles highlight *P. tremula* individuals (*q*<0.05) and *P. alba* individuals (q>0.95), respectively, while the grey rectangle indicates hybrids. 95% confidence intervals are too small to be depicted. (**b**) Local ancestries along the chromosomes of 28 exemplary individuals (each row is an individual), representing the range of variation of *q*. Confidence in ancestry estimates is shown by shades from white (unknown ancestry) to blue (*P. alba* ancestry), orange (*P. tremula* ancestry) or grey (heterospecific ancestry). See Fig. S3 for the results of all individuals.

### Local ancestry inference

We estimated local ancestry using RASPberry (Wegmann *et al*., 2011) based on 32,413 SNVs passing filters and under a model with five generations since admixture, ancestral recombination rates of 500, and a miscopying rate of 0.06, which had the highest likelihood. Local ancestry analysis revealed a genomic mosaic of homospecific ancestry segments derived from *P. alba* and *P. tremula* and segments with heterospecific ancestry (Fig. **1b**). As expected from genome-wide ancestries, we observed more *alba*-like than *tremula*-like hybrids in our sample set (Fig. **S3**).

### Admixture LD

Successful mapping in any association study depends on the extent of LD between sites (Remington *et al*., 2001; Stracke *et al*., 2007). LD in our data displayed spatial decay patterns along chromosomes suitable for phenotype mapping: adjacent loci showed strong LD of ancestry state, which decayed gradually with physical distance (Figs. **2**, **S4**, **S5**).

**Fig. 2.**
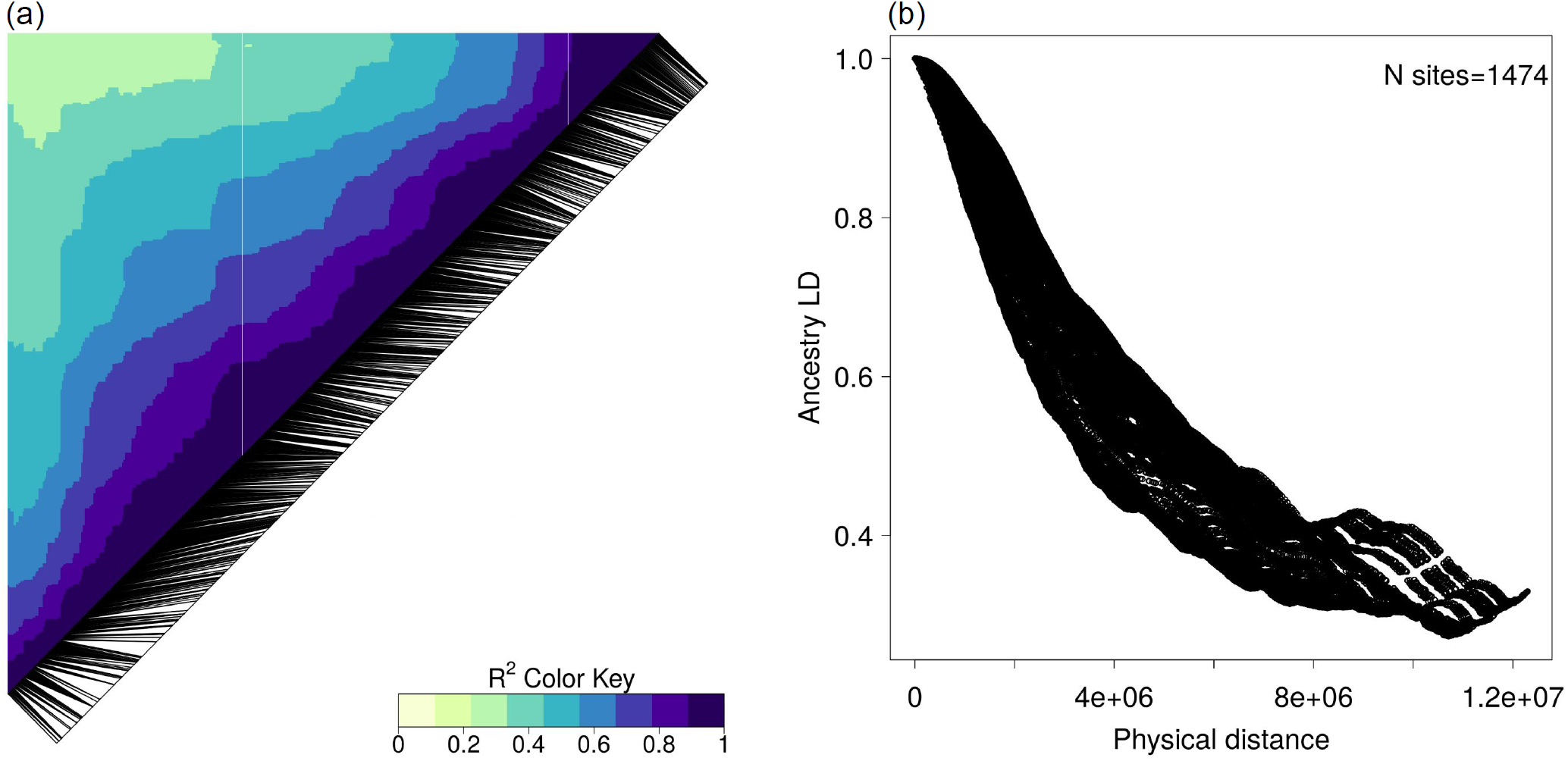
AdmixtureLD on chromosome 9. (**a**) Black lines indicate the positions of analyzed loci along the chromosome and darker blue shades represent stronger LD. (**b**) LD decay as a function of physical distance along the chromosome. *N sites* indicates the number of loci analyzed on this chromosome. Results for the remaining chromosomes were very similar (Figs. S4, S5).

### Phenotypic differentiation

Traits in our GWAS showed variable levels of differentiation between *P. alba* and *P. tremula.* Two example traits are shown in Fig. **3**: C12, which showed strong divergence between *P. alba* and *P. tremula* and intermediate values in hybrids, and C34, which exhibited similar abundance in all genotypic classes. For box plots summarizing patterns of variation for all traits see Figs. **S6**, **S7**. For some traits, we detected transgressive phenotypes in hybrids (Fig. **3b,d**; e.g. C34). The proportion of phenotypic variance explained by genome-wide ancestry (*q*) followed a heterogeneous pattern (Fig. **4a**), broadly mirroring patterns of intra- and interspecific variability for the studied traits (Fig. **S7**).

**Fig. 3.**
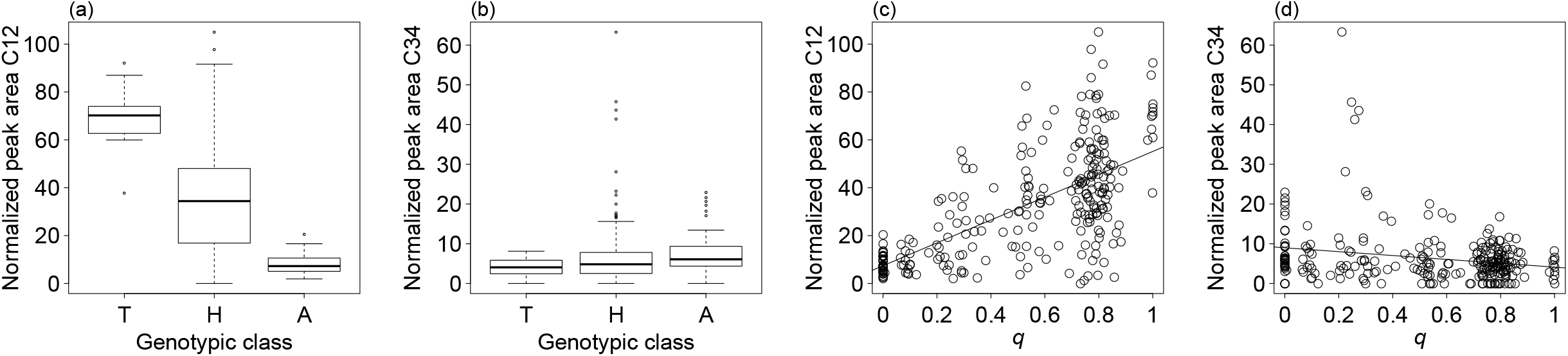
(**a**) and (**b**) show levels of differentiation between *P. alba*, *P. tremula* and their hybrids for two phytochemical traits (C12 and C34, respectively). T, *P. tremula*(*q* <0.05); H, hybrid seedlings with 0.05≤*q*≤0.95; A, *P.alba* (*q*>0.95). (**c**) and (**d**) depict the relationship between genome-wide ancestry (*q*, x axis) and the two phytochemical traits. *P. tremula*-like individuals are on the left, where *q* <0.05, while *P. alba*-like individuals are on the right, where *q* >0.95. Hybrid seedlings exhibit intermediate values of q. Linear regression lines are shown as visual guides only and are not intended to suggest that a linear regression function represents the best fit to the data.

**Fig. 4.**
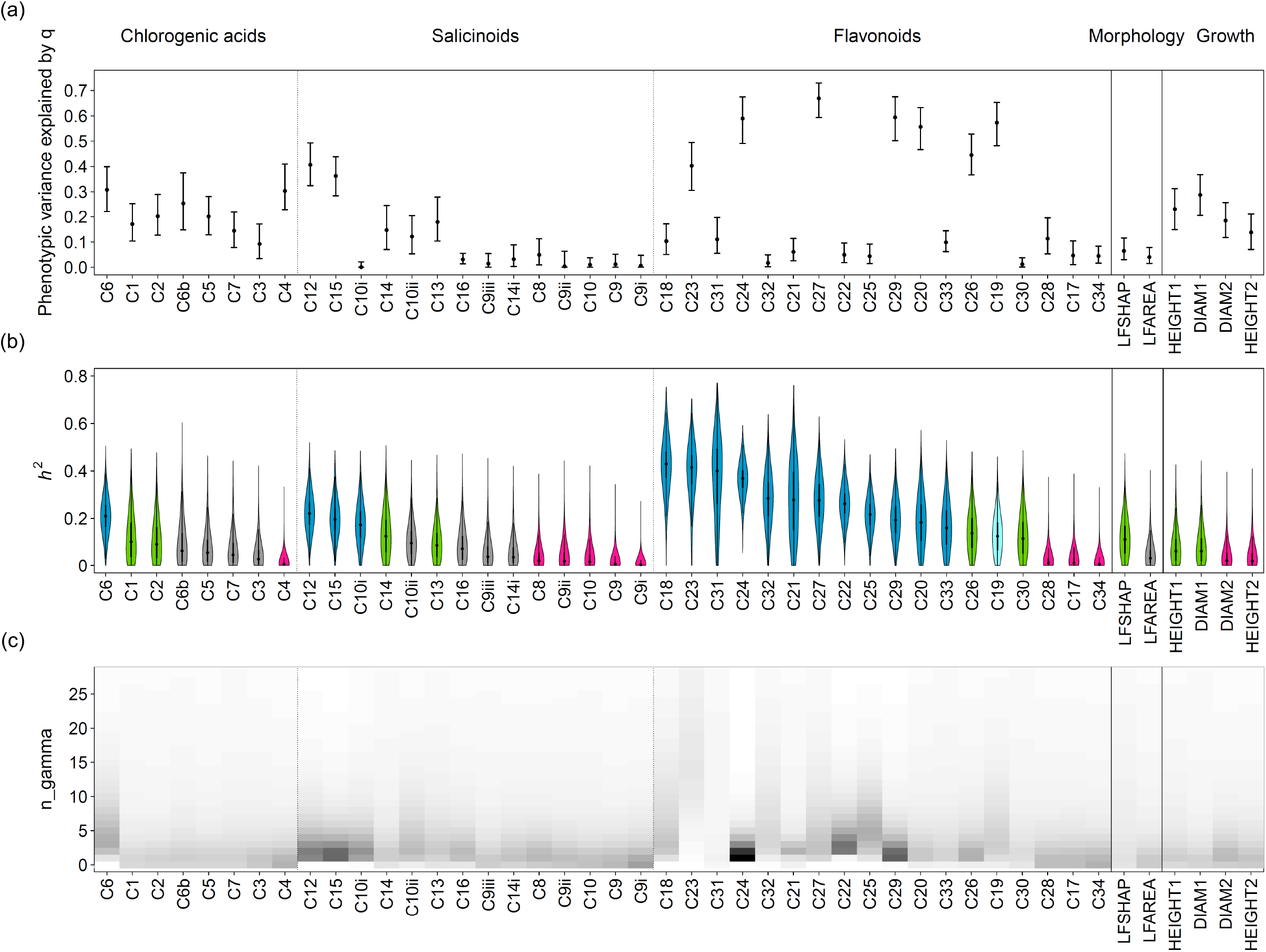
Results for all traits analyzed in this GWAS, grouped based on functional similarities among traits (phytochemistry including chlorogenic acids, salicinoids, and flavonoids; morphology; growth) and ordered according to the decreasing median of narrow-sense heritability *h^2^*. (**a**) Proportion of phenotypic variance explained by *q*. Bars indicate 95% confidence intervals. (**b**) Violin plots showing the posterior distributions of *h^2^* of traits assigned to the first class of genomic architecture (blue), to the second class (green), to the third class (pink), and of traits which could not be assigned to any class (gray). C19 is shown in light blue, since it only barely missed our threshold on *h^2^* to be included in the first class and showed a sharp PIP peak (see text). (**c**) Heatmap for the values of *n_gamma*. Darker shades indicate higher number of occurrences for the corresponding value of *n_gamma* in the posterior distribution.

### Admixture mapping

Parameter estimates obtained with the BSLMM implemented in GEMMA revealed a continuum of genomic architectures (Figs. **4b**, **S8**, Table S1). For further analysis, we grouped them into three main classes (Table 2):

**Table 2.** Traits assigned to each class of genomic architecture, as suggested by hyperparameter posterior distributions from GEMMA.

| Trait class | Inclusion criteria | Traits<sup>a</sup> |
| --- | --- | --- |
| First | $h^2 \geq 0.01$ with >95% probability | C6, C10i, C12, C15, C18, C20, C21, C22, C23, C24, C25, C27, C29, C31, C32, C33, C19<sup>b</sup> |
| Second | $h^2 \geq 0.01$ with <95% probability, but $PVE \geq 0.05$ with >97% probability | C1, C2, C13, C14, C26, C30, HEIGHT1, DIAM1, LFSHAP |
| Third | $h^2 < 0.01$ with $\geq 30\%$ probability | C4, C8, C9, C9i, C9ii, C10, C17, C28, C34, HEIGHT2, DIAM2 |
| Not assigned | --- | C3, C5, C6b, C7, C9iii, C10ii, C14i, C16, LFAREA |

<sup>a</sup>Trait abbreviations as in Table 1
<sup>b</sup> This trait does not strictly satisfy the requirements for the first class, but was included because it showed a strong signal for heritability ( $h^2 \geq 0.01$ with >93% probability) and a sharp Posterior Inclusion Probability (PIP) peak (see text).

The *first class* included traits with strong evidence for heritability and with both the genomic background and loci with measurable effect contributing to the phenotypic variation. This class of loci with strong evidence for a genetic role in explaining phenotypic variation included 16 phytochemical traits with *h^2^* ≥0.01 with at least 95% probability (Table 2; Figs. **4b**, **5a**). These were the phytochemical traits showing the highest values of median *h^2^* and highest probability of *n_gamma* >0 (Fig. **4b,c**). An additional trait (C19) was considered part of this class, although it did not strictly satisfy the threshold on *h^2^* (see below).

The *second class* corresponds to traits for which only the genomic background appears to play a role in explaining the phenotype, while the actual contribution of individual loci with measurable effect on the phenotype is less clear. This group encompasses six phytochemical traits, the growth traits DIAM1 and HEIGHT1 and the morphological trait LFSHAP (Table 2; Figs. **4b**, **5b**), which do not meet the heritability threshold outlined above, but for which *PVE* was ≥0.05 with >97% probability.

The *third class* included traits for which both the genomic background and the variation explained by loci with measurable effect were not significantly different from zero, thus causing *h^2^* to approach zero. The most evident cases for this scenario were phytochemical traits C4 (Fig. **5c**), C9, C9i and C34, for which *h^2^*<0.01 with >60% probability and *h^2^* <0.05 with >90% probability. The probability of *n_gamma* =0 was the highest for these traits. In total, nine phytochemical traits and the growth traits DIAM2 and HEIGHT2 were identified to be in this class (Tables 2, S1).

**Fig. 5.**
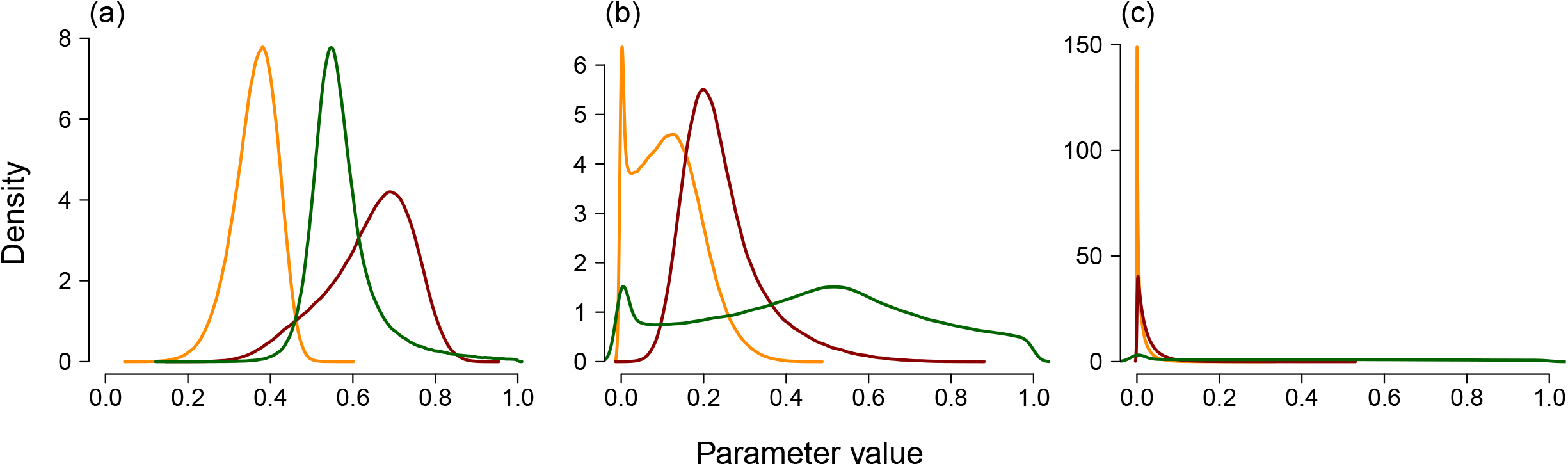
Posterior distributions of *PVE* (red), *PGE* (green) and *h^2^* (yellow) for an exemplary trait of each class: (a) C24 for the first class, (b) LFSHAP for the second class, and (c) C4 for the third class.

For the remaining traits (seven phytochemical traits and LFAREA; Table 2), the posterior distributions of *h^2^* did not allow us to obtain clear insights concerning their genomic architecture. It was therefore not possible to assign them to a specific class.

### Analysis of focal traits with finite genomic architecture

We identified the genomic regions with sparse effects for the 16 traits in the first class of genomic architectures. All of these were phytochemical traits, i.e. secondary metabolite compounds: one chlorogenic acid (C6, *5-coumaroyl quinic acid*), three salicinoids (C10i, *acetyl-salicortin isomer 1*; C12, *HCH-salicortin*; C15, *HCH-tremulacin*) and 12 flavonoids (C18, *quercetin-rutinoside-pentose*; C20, *quercetin-hexose-pentose*; C21, *kaempferol-rutinoside-pentose;* C22, *isorhamnetin-rutinoside-pentose;* C23, *quercetin-3-O-rutinoside*; C24, *quercetin-3-O-glucuronide*; C25, *quercetin-3-O-glucoside*; C27, *isorhamnetin-3-O-rutinoside*; C29, *kaempferol-glycuronide*; C31, *isorhamnetin-glycoside*; C32, *isorhamnetin-glycuronide*; C33, *isorhamnetin-acetyl-hexose).* As mentioned above, we investigated an additional trait as part of this set (C19, *quercetin-glucuronide-pentose*), since it exhibited a genomic window with PIP ≥0.4 and only barely missed our threshold on *h^2^* with a posterior probability >93%.

Eleven traits with finite effects were associated with a single genomic window with PIP ≥0.4 per trait, while traits C18, C22, C23, C31, C32 exhibited two or three windows above this threshold. C33 had no genomic window with PIP ≥0.4, despite satisfying the requirements regarding *h^2^* and *n_gamma*. Since the phytochemical compounds underlying traits C19, C29 and C32 were not produced by >10% of individuals, we also conducted mapping on the presence or absence of these compounds (binary analysis, Notes S1). This analysis also revealed identifiable sparse effects, but only for C29 did the signal reside in the same genomic window as in the quantitative analysis. For traits C19 and C32, in contrast, the identified windows did not overlap, suggesting that different variants are responsible for down-regulating or inhibiting the pathway leading to these compounds.

Windows of special interest were located on chromosomes 1, 3, 6, 11, 12, 13, 15 and 18 (Fig. **6**; Notes S2; Table S2). Several windows showed PIP peaks for more than one trait: this was the case for two windows on each of the chromosomes 11, 12 and 15. Particularly interesting is the window between 3 and 3.5 Mb on chromosome 12, which appears to be involved in explaining six different phytochemical traits, all belonging to the flavonoid branch of the phenylpropanoid pathway. Results obtained with alternative modeling options in GEMMA (Methods S3) corroborated those obtained with our primary approach (Notes S1).

**Fig. 6.**
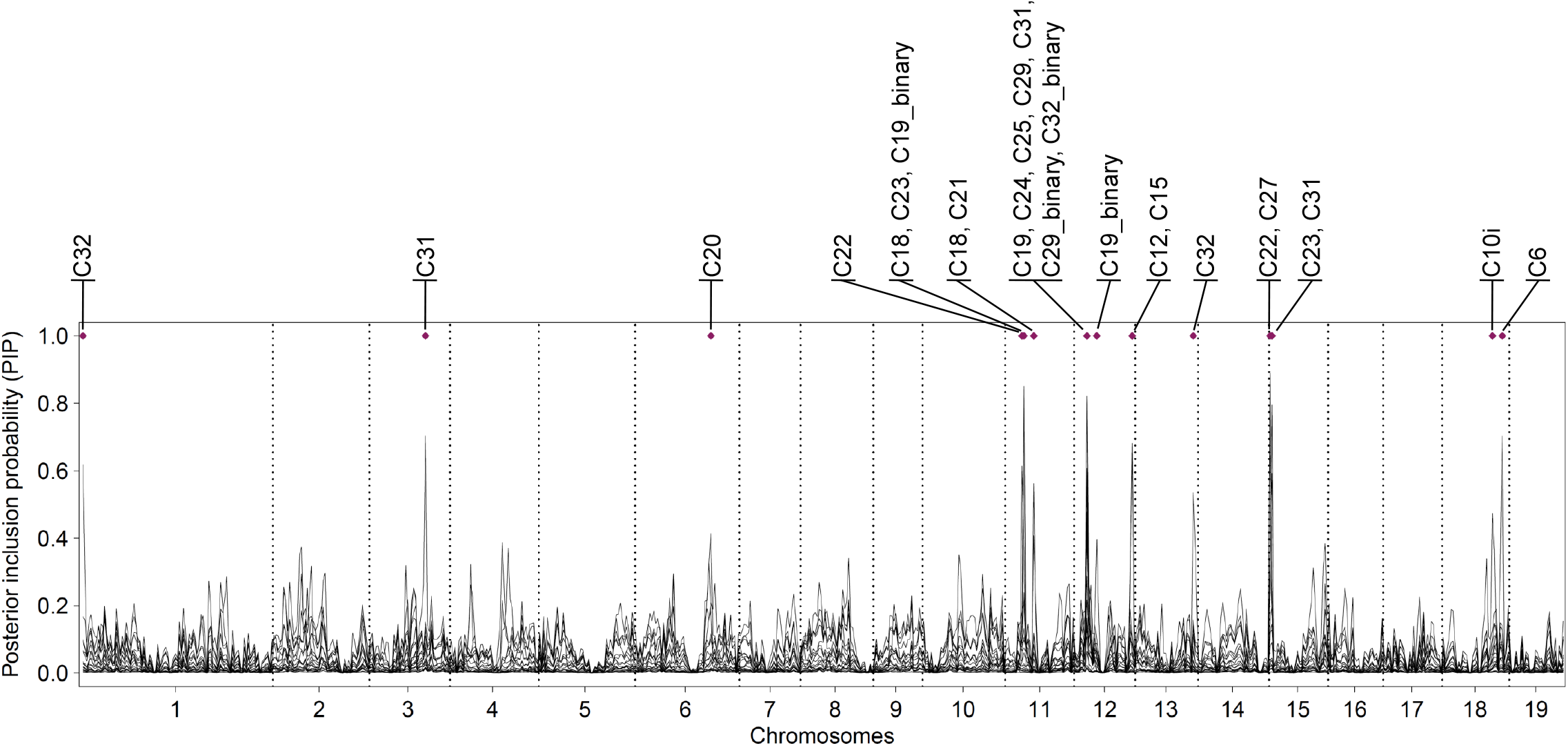
Values of Posterior Inclusion Probabilities (PIPs) per 0.5 Mb windows for all selected traits. Windows with PIP ≥0.4 are marked with a violet diamond and the traits where this threshold is exceeded are indicated.

### Candidate genes in windows with high PIP

Within the windows of interest (PIP ≥0.4), we identified several candidate genes potentially responsible for the significant associations between specific genomic regions and phenotypic traits. Most conspicuously, these windows contained genes encoding *MYB-type transcription factors* known to function in combination in plants (Höll *et al*., 2013; Liu *et al*., 2014; *P. trichocarpa* gene models Potri.001G005100, Potri.013G149100, and Potri.013G149200); several *UDP-glycosyl transferases*, i.e. proteins able to transfer sugar moieties with important roles in phenylpropanoid compound biosynthesis (Aksamit-Stachurska *et al*., 2008; Babst *et al*., 2014; Caseys *et al*., 2015; e.g. Potri.011G060300 and Potri.011G061000); a *glycosyl hydrolase* involved in the biosynthetic flow by reducing the complexity of sugars of compounds (Potri.015G010100); COMT1, a *methyl transferase* that transforms quercetins into isorhamnetins (Potri.015G003100) and CHS, a *chalcone synthase* (Potri.012G138800). More information on candidate genes found in genomic windows with high PIP is provided in Notes S2 and Table S2, and the most plausible candidate genes are discussed below.

## Discussion

Early genetic mapping studies have often pointed to potentially simple, sparse genetic architectures of phenotypic traits in wild and domesticated species (reviewed by e.g. Coyne & Orr, 2004), but we now know that adaptive traits are often polygenic (e.g. Pritchard *et al*., 2010; Rockman, 2012; Evans *et al*., 2014; Pasaniuc & Price, 2017) or possibly even “omnigenic” (Boyle *et al*., 2017). Given sufficient power and suitable analytical tools, a reasonable expectation for trait architectures uncovered by mapping studies may thus be no or only very subtle genetic effects, unless (1) multiple small-effect mutations or pleiotropic effects accumulate in the same hotspot of phenotypic evolution (Martin & Orgogozo, 2013), (2) the traits are very tightly coupled with (or the direct products of) the underlying biochemical pathways (e.g. Boeckler *et al*., 2011), or (3) the focus is on traits segregating between highly divergent populations, for which architectures may differ from classical within-species expectations (Rieseberg & Buerkle, 2002; Lexer *et al*., 2005). In this study, we have investigated the genomic architecture of phenotypic trait differences between two ecologically divergent forest tree species (*Populus alba* and *P. tremula*) by applying GWAS to an admixed population.

### Variation available for genetic mapping in admixed populations

Admixture mapping studies require sufficient genetic and phenotypic variation to uncover genetic associations and trait architectures (Briscoe *et al*., 1994; reviewed by Buerkle & Lexer, 2008). Our analysis of genomic variation confirmed that these two poplar species, besides their ecological divergence, are also strongly divergent genetically at many loci (mean F_ST_ 0.392), in line with previous estimates (Lexer *et al*., 2010; Stölting *et al*., 2013; Christe *et al*., 2017). This resulted in favorable conditions for estimating local ancestry (Christe *et al*., 2016) and thus for mapping. Divergence at the genetic level was reflected by pronounced differentiation at the phenotypic level for a range of characters (Figs. **3**, **S6**, **S7**), with several traits showing strong differences between the parental species, especially in the case of phytochemical traits. Hybrids showed intermediate or parental-like values for most traits. Nevertheless, many recombinant hybrids showed phenotypic values falling outside the range of variation of the parental species (Figs. **3b,d**, **S6**, **S7**) and are thus examples of transgressive segregation (Rieseberg *et al*., 2003; Dittrich-Reed & Fitzpatrick, 2013).

### Traits with high, medium, or low heritability

Our GWAS identified genomic architectures that fall roughly into three main classes. The first class consisted of traits with relatively high heritability *h^2^* and for which a finite set of genomic regions contribute to the phenotype. All these were phytochemical traits, a finding consistent with the notion that finite genetic effects are more easily detected for traits tightly coupled with the underlying pathways.

The second class included traits for which phenotypic variation is explained by genetic effects as detected by *PVE* captured by our kinship (=genomic similarity) matrix, but no localized association was identified (*PGE* about zero). One likely reason why the genomic similarity matrix explains phenotypic variation is that the trait is heritable but highly polygenic, as was previously reported for growth-related traits (Wood *et al*., 2014; Tsai *et al*., 2015), also in the case of *Populus* species (Du *et al*., 2016).

The third class consisted of traits that do not appear to be heritable. Many of these are phytochemical defense traits against herbivores, which may be predominantly influenced by environmental factors (Abreu *et al*., 2011; Boeckler *et al*., 2011). However, some of these traits did show considerable phenotypic differences between the species, also in our common garden setting. The potential lack of a heritable signal could therefore also indicate a lack of power of our admixture mapping approach. Indeed, many causal variants were likely missed by our reduced representation sequencing experiment, and also not well tagged due to generally very low LD in *Populus* (few hundred base pairs according to Ingvarsson, 2008; Marroni *et al*., 2011; but see Olson *et al*., 2010; Slavov *et al*., 2012). To mitigate this issue, however, we chose to conduct mapping on local ancestry, which exhibits long-range LD among the early generation hybrids used here. A more likely cause for the inferred trait architectures is thus that some of these traits are highly polygenic, and a failure to detect significant heritability for such traits might be due to a lack of power associated with admixture mapping.

### Low heritability of growth-related traits

Heritability estimates were conspicuously low for growth-related traits, despite controlling for potential covariates. A lack of heritable variation for growth traits was previously observed in poplar and willow species (Orians *et al*., 2003; Du *et al*., 2016). The *PVE* estimates we observed are likely mainly due to the phenotypic variance that can be explained by the genomic ancestry similarity matrix (Zhou *et al*., 2013) as discussed earlier. Nevertheless, other explanations have to be taken into account as well.

### Pleiotropic effects among phytochemical traits

Out of the 14 genomic windows exhibiting high values of PIP, six showed significant association with more than one phytochemical trait. This suggests that loci controlling different compounds are in linkage in the same window, or that the same loci are responsible for several traits, i.e. that they have pleiotropic effects. One indication suggesting pleiotropic effects is that the traits associated with the same window always belong to the same branch of the phenylpropanoid pathway (five windows associated with several flavonoids and one window associated with two salicinoids). These windows could host enzymes acting upstream in a specific pathway branch, thus affecting several downstream steps and compounds (Cork & Purugganan, 2004).

### Candidate genes associated with phenylpropanoid compounds

We identified several noteworthy candidate genes in PIP peaks for flavonoid traits that code for glycosyl transferases. These enzymes act in glycosylation (i.e. conjugation to a sugar moiety), one of the most widespread modifications of plant secondary metabolites (Gachon *et al*., 2005), effectively modifying solubility, stability, and reactivity of compounds (Aksamit-Stachurska *et al*., 2008). Their association with genes within PIP peaks for flavonoid traits is consistent with their previously inferred involvement in transgressive expression of phytochemical traits in an overlapping sample of poplar hybrids (Caseys *et al*., 2015). Among other noteworthy candidate genes (Notes S2), a chalcone-synthase (CHS) within a PIP peak for HCH-salicinoids (two highly toxic compounds) is of particular interest: this gene, currently assigned to the flavonoid pathway, may instead be active in the largely unknown salicinoid pathway. We hypothesize that the polyketide synthase activity may directly act on benzoyl-CoA, which has recently been put forward as a likely precursor of this entire group of compounds (Notes S2).

The genomic windows with high PIP also yielded a short list of candidate genes for the biosynthesis of the studied phenylpropanoid compounds and its regulation (Table S2). The flavonoid isorhamnetin-glycuronide (C32), for example, was significantly associated with two windows hosting three transcription factors of the MYB family previously shown to regulate the phenylpropanoid pathway (Sablowski *et al*., 1994; Liu *et al*., 2015). The candidate genes MYB14 and MYB15 were found to interact in plants to stimulate the production of stilbenes, a group of phenylpropanoid compounds produced in response to biotic and abiotic stress (Höll *et al*., 2013; Duan *et al*., 2016). MYB15 also confers improved tolerance to drought and salt stress (Ding *et al*., 2009), negatively regulates the expression of CBFs (genes expressed in response to cold conditions; Chinnusamy *et al*., 2007), and regulates defense-induced lignification and basal immunity in *A. thaliana* (Chezem *et al*., 2017). Finally, the transcription factor MYB5, which is under positive selection in *P. tremula* (Christe *et al*., 2017), interacts physically with MYB14 and activates the promoter of enzymes related to the biosynthesis of pro-anthocyanidins (Liu *et al*., 2014), a major class of flavonoids responsible for color in various plant organs.

These findings are remarkable from the perspective of reproductive isolation between divergent species and in particular the breakdown of hybrid fitness in *P. alba* and *P. tremula* hybrids (Christe *et al*., 2016): postzygotic reproductive barriers could originate from the disruption of coadapted gene complexes, when proper interactions between gene products cannot take place and consequently, hybrids show a non-functional phenotype (Ortíz-Barrientos *et al*., 2007; Livingstone *et al*., 2012; Lindtke & Buerkle, 2015). These interacting MYB transcription factors might represent an example of this mechanism at work: their involvement in plant defense could cause adverse effects in plants carrying incompatible genotypes, thus affecting plant survival and performance at early life stages when selection in trees tends to be strong (Petit & Hampe, 2006).

## Acknowledgements

We are grateful to Kai N. Stölting, Camille Christe, Margot Paris, Dorothea Lindtke, Berthold Heinze and members of the Wegmann lab for their help and advice. We thank also Thelma Barbará, Andrew Brown, David Frey, Alberto Spada, Angela Cicatelli, Franscesco Guarino, and the botanical garden teams at Fribourg and Salerno. This work was supported by grant 31003A_149306 of the Swiss National Foundation to CL.

## Author Contribution

CL and LB conceived the study; CL provided funding; LB, CC and SC collected genetic and phenotypic data; LB and CAB performed the analyses; CC collected information and provided valuable insights about candidate genes; CAB, DW and CL supervised the study; LB, DW and CL wrote the manuscript with input and revisions from all co-authors.

## Supporting Information

**Fig. S1** Posterior inclusion probabilities in genomic windows of different size

**Fig. S2** Kinship matrix calculated by GEMMA

**Fig. S3** Local ancestries for all analyzed seedlings

**Fig. S4** Admixture linkage disequilibrium in all chromosomes

**Fig. S5** Decay of admixture linkage disequilibrium along all chromosomes

**Fig. S6** Levels of phenotypic differentiation between *P. tremula*, hybrids and *P. alba* for all traits

**Fig. S7** Relationship between genome-wide ancestry and phenotype for all traits

**Fig. S8** Posterior distributions of *PVE, PGE* and heritability for all phenotypic traits

**Table S1** Probabilities from posterior distributions of heritability, *PVE, PGE* and *n_gamma*

**Table S2** Candidate genes in genomic windows with high posterior inclusion probability

**Methods S1** RAD-seq data processing, reference-mapping and variant calling

**Methods S2** Inference of local and genome-wide ancestry

**Methods S3** Rationale for choice of plant traits measured in this study

**Methods S4** Admixture mapping with GEMMA: model choice and validation

**Notes S1** Genomic windows highlighted by alternative modeling approaches in GEMMA

**Notes S2** Additional information on candidate genes

